# Translation-associated mutational U-pressure in the first ORF of SARS-CoV-2 and other coronaviruses

**DOI:** 10.1101/2020.05.05.078238

**Authors:** Khrustalev Vladislav Victorovich, Giri Rajanish, Khrustaleva Tatyana Aleksandrovna, Kapuganti Shivani Krishna, Stojarov Aleksander Nicolaevich, Poboinev Victor Vitoldovich

**Author notes:** Correspondence to: Dr. Vladislav V Khrustalev, Department of General Chemistry, Belarusian State Medical University, Dzerzinskogo 83, Minsk, Belarus.

## Abstract

Within four months of the ongoing COVID-19 pandemic caused by SARS-CoV-2, more than 250 nucleotide mutations have been detected in the ORF1 of the virus isolated from different parts of the globe. These observations open up an obvious question about the rate and direction of mutational pressure for further vaccine and therapeutics designing. In this study, we did a comparative analysis of ORF1a and ORF1b by using the first isolate (Wuhan strain) as the parent sequence. We observed that most of the nucleotide mutations are C to U transitions. The rate of synonymous C to U transitions is significantly higher than the rate of nonsynonymous ones, indicating negative selection on amino acid substitutions. Further, trends in nucleotide usage bias have been investigated in 49 coronaviruses species. A strong bias in nucleotide usage in fourfold degenerated sites towards uracil residues is seen in ORF1 of all the studied coronaviruses. A more substantial mutational U pressure is observed in ORF1a than in ORF1b owing to the translation of ORF1ab via programmed ribosomal frameshifting. Unlike other nucleotide mutations, mutational U pressure caused by cytosine deamination, mostly occurring in the RNA-plus strand, cannot be corrected by the proof-reading machinery of coronaviruses. The knowledge generated on the direction of mutational pressure during translation of viral RNA-plus strands has implications for vaccine and nucleoside analogue development for treating covid-19 and other coronavirus infections.

## Introduction

The current COVID-19 pandemic caused by SARS-CoV-2 has claimed more than 0.25 Million human lives with nearly 3.5 million reported infections globally and still this number is increasing day by day.^1^ The virus was first detected in a wet market in Wuhan, China in December 2019. It bears a close similarity to bat coronaviruses, and similar viral particles have been detected in pangolin. This led to the speculation that the virus must have originated in bats while pangolins must have been intermediate hosts. Since then, the virus has spread all over the globe affecting humans of all races. The infections are characterized by sore throat, fever, cough, body pains, breathlessness, severe pneumonia, and death due to multi-organ failure involving kidneys and lungs.^2,3^

Coronaviruses have an exceptionally large genome of around 30kb. A huge part of the genome, at the 5’ end, around 20 kb, harbors a replicase gene that codes for around 16 non-structural proteins (nsps).^4^ The rest of the genome, near the 3’ end, codes for the structural (spike, envelope, membrane, nucleocapsid) and several accessory proteins.^5^ The replicase gene (ORF1) includes two subunits, ORF1a and ORF1b expressing the polyproteins pp1a and pp1ab, respectively. A slippery sequence and ribosomal frameshifting caused by an RNA pseudoknot ensure the translation of both polyproteins from the same genome.^6,7^ Pp1a has nsps 1-11 whereas pp1ab has nsps 1-16.^8^ The non-structural proteins form the viral replicase complexes. Here both genomic and subgenomic viral RNAs get synthesized through negative-strand intermediates. Sub genomic RNAs code for structural and accessory proteins.

Variations in viral sequences have an important role in viral propagation and pathogenesis. They may help change the viral phenotype allowing it to evade host immune system and also acquire drug resistance. These variations may arise from copying errors during genome replication.^9^ RNA viruses have low replicative fidelity; hence are more prone to mutations than DNA viruses. Around 10^−4^ to 10^−6^ errors per nucleotide are seen in RNA viruses, which is equivalent to approximately one mutation per genome per replication cycle. This allows the viruses to maintain mutations that are beneficial for them in adapting to new environments.^10^ Sometimes, a specific nucleotide mutation occurs more frequently, which is called directional mutational pressure. Mutational pressure changes the nucleotide composition of the genome irreversibly and may be caused due to an error-prone polymerase, RNA editing, oxidative damage. Moreover, the direction of mutational pressure may not be similar along the entire length of the genome.^11^ Coronaviruses are known as slowly mutating RNA viruses since they perform proof-reading during replication^12^.

Cytosine to Uracil transitions can occur spontaneously through oxidative damage by free radicals or enzymatically through the action of apolipoprotein B mRNA editing catalytic subunit (APOBEC) family of cytidine deaminases. RNA-editing enzymes from APOBEC family bind single-stranded viral RNA and deaminate cytosine residues.^13,14^ Single nucleotide transitions mostly result in synonymous mutations though synonymous codons, i.e., codons that code for same amino acid, don’t occur equally in different organisms; these organisms are said to have a codon usage bias. Fourfold degenerate codons can tolerate any nucleotide substitution at the wobble or the third position, which doesn’t change the amino acid sequence. But this changed nucleotide in the RNA may affect the RNA structure, RNA interference, and codon usage bias.^11^

Since its first appearance in December, 2019, four hundred full-length sequences of ORF1 of SARS-CoV-2 have become available on GenBank. In this study, a comparative analysis of ORF1a and ORF1b has been carried out, taking the first isolate as the parent sequence. Further, trends in nucleotide usage bias have also been investigated in various coronaviruses. There are no approved drugs or vaccines for human coronaviruses yet. With alarmingly increasing damage caused by COVID-19 to human lives and global economy, there is a pressing need for its prevention and treatment strategies through vaccines or potential drugs. This knowledge on direction of mutational pressure may help design more effective vaccines and nucleoside analogues.

## Materials and Methods

### Data/ Nucleotide sequence retrieval

The nucleotide sequences of ORF1 from genomes of 49 coronaviruses were retrieved from GenBank. These include Alphacoronaviruses, Betacoronaviruses, Gammacoronaviruses, and Deltacoronaviruses (Table 3) including coronaviruses that caused an epidemic in 2003 (SARS-CoV), in 2013 (MERS-CoV), and an ongoing pandemic that started in December 2019 (SARS-CoV-2).

**Table 1.** Rates of nucleotide mutations (in substitution per site with a corresponding nucleotide) in two parts of SARS-CoV-2 ORF1: before (ORF1a) and after (ORF1b) the ribosome slippage sequence. Significant changes between opposite types of mutations are shown by bold font, significant changes in rate of the same type of mutation between two parts of ORF1 are shown in underlined font.

| Type of mutation: | C→U | U→C | A→G | G→A | C→A | A→C |
| --- | --- | --- | --- | --- | --- | --- |
| ORF1a | **<u>0.0356223</u>** | **0.005379** | 0.004306 | <u>0.007928</u> | 0.001288 | 0.000760 |
| ORF1b | **<u>0.0212164</u>** | **0.004971** | 0.003633 | <u>0.001900</u> | 0.001414 | 0.000404 |
| Type of mutation: | G→U | U→G | A→U | U→A | G→C | C→G |
| ORF1a | **0.0049075** | **0.000000** | 0.000760 | 0.000935 | 0.000378 | 0.000429 |
| ORF1b | **0.0082331** | **0.000382** | 0.000404 | 0.001147 | 0.000633 | 0.000707 |

**Table 2.** Description of C to U transitions in two parts of SARS-CoV-2 ORF1: before (ORF1a) and after (ORF1b) the ribosome slippage sequence. Significant changes between rates of synonymous and nonsynonymous mutations are shown by bold font, significant changes in rate of the same type of mutation between two parts of ORF1 are shown in underlined font.

|  | ORF1a |  | ORF1b |  |
| --- | --- | --- | --- | --- |
|  | synonymous | nonsynonymous | synonymous | nonsynonymous |
| number of sites with C to U mutation | 32 | 51 | 17 | 13 |
| number of available sites | 630 | 1700 | 403 | 1011 |
| rate of mutations, substitution per site | **0.050794** | **<u>0.03000</u>** | **0.042184** | **<u>0.012859</u>** |

**Table 3.** Average values of nucleotide content in fourfold degenerated sites for two parts of ORF1 (before (ORF1a) and after (ORF1b) the ribosome slippage sequence) for 49 species of coronaviruses.

| Species of viruses | ORF1a |  |  |  | ORF1b |  |  |  |
| --- | --- | --- | --- | --- | --- | --- | --- | --- |
| **Betacoronaviruses** | A4f | U4f | G4f | C4f | A4f | U4f | G4f | C4f |
| SARS-CoV-2 | 0.282559 | 0.536080 | 0.058141 | 0.123220 | 0.314966 | 0.489943 | 0.065602 | 0.129489 |
| Bat coronavirus RaTG13 | 0.279333 | 0.538640 | 0.057880 | 0.124147 | 0.317106 | 0.492413 | 0.063180 | 0.127301 |
| SARS type 1 | 0.267316 | 0.492572 | 0.080709 | 0.159403 | 0.326612 | 0.429282 | 0.084896 | 0.159210 |
| Betacoronavirus England 1 | 0.223760 | 0.492141 | 0.096139 | 0.187959 | 0.222440 | 0.505384 | 0.087213 | 0.184963 |
| MERS | 0.223758 | 0.489458 | 0.097238 | 0.189546 | 0.221945 | 0.506858 | 0.086123 | 0.185074 |
| Bat coronavirus HKU4-1 | 0.243970 | 0.549858 | 0.086605 | 0.119567 | 0.247374 | 0.527550 | 0.098643 | 0.126433 |
| Bat coronavirus HKU5-1 | 0.219651 | 0.430946 | 0.126756 | 0.222647 | 0.233393 | 0.428189 | 0.137513 | 0.200905 |
| Bat coronavirus HKU9-1 | 0.205231 | 0.517617 | 0.143563 | 0.133589 | 0.248432 | 0.471643 | 0.135637 | 0.144288 |
| Bat Hp-betacoronavirus Zhejiang 2013 | 0.259312 | 0.449471 | 0.127087 | 0.164130 | 0.280761 | 0.428425 | 0.109822 | 0.180992 |
| Bovine coronavirus TCG-19 | 0.210180 | 0.594721 | 0.101922 | 0.093176 | 0.229055 | 0.512969 | 0.110493 | 0.147483 |
| China rattus coronavirus HKU24 | 0.220346 | 0.470419 | 0.147173 | 0.162062 | 0.244320 | 0.445679 | 0.136765 | 0.173236 |
| Betacoronavirus Erinaceus | 0.294150 | 0.482614 | 0.109619 | 0.113616 | 0.303022 | 0.478903 | 0.082157 | 0.135918 |
| Human coronavirus HKU1 | 0.207154 | 0.701287 | 0.037482 | 0.054077 | 0.217268 | 0.643761 | 0.049733 | 0.089239 |
| Human coronavirus OC43 | 0.212476 | 0.590722 | 0.09732 | 0.099481 | 0.237404 | 0.514365 | 0.101690 | 0.146541 |
| Mouse hepatitis virus | 0.188042 | 0.448827 | 0.166824 | 0.196306 | 0.216405 | 0.421702 | 0.166198 | 0.195695 |
| Rabbit coronavirus HKU14 | 0.196470 | 0.590026 | 0.112055 | 0.101449 | 0.252936 | 0.497622 | 0.100746 | 0.148696 |
| Rat coronavirus Parker | 0.186994 | 0.459502 | 0.167466 | 0.186037 | 0.214727 | 0.438168 | 0.157116 | 0.189988 |
| Rousettus bat coronavirus | 0.240135 | 0.400558 | 0.175626 | 0.183681 | 0.262846 | 0.396113 | 0.177551 | 0.163490 |
| **Alphacoronaviruses** | A4f | U4f | G4f | C4f | A4f | U4f | G4f | C4f |
| Bat coronavirus 1A | 0.197439 | 0.592162 | 0.064526 | 0.145873 | 0.201381 | 0.610753 | 0.061812 | 0.126054 |
| Bat coronavirus CDPHE15USA2006 | 0.211126 | 0.559267 | 0.087088 | 0.142519 | 0.192782 | 0.557644 | 0.087228 | 0.162345 |
| Bat coronavirus HKU2 | 0.166628 | 0.652947 | 0.062911 | 0.117514 | 0.199027 | 0.584848 | 0.083031 | 0.133093 |
| BtMr-AlphaCoV SAX2011 | 0.209349 | 0.556779 | 0.097402 | 0.136470 | 0.238494 | 0.507276 | 0.081311 | 0.172918 |
| BtNv-AlphaCoVCS2013 | 0.161644 | 0.585640 | 0.111608 | 0.141108 | 0.177368 | 0.559410 | 0.106309 | 0.156914 |
| BtRf-AlphaCoV HuB2013 | 0.245029 | 0.558607 | 0.084154 | 0.112210 | 0.261599 | 0.490377 | 0.101753 | 0.146271 |
| BtRf-AlphaCoVYN2012 | 0.180421 | 0.662049 | 0.054639 | 0.102891 | 0.205510 | 0.596575 | 0.06731 | 0.130605 |
| Camel alphacoronavirus | 0.208933 | 0.593785 | 0.076436 | 0.120846 | 0.235353 | 0.546779 | 0.088768 | 0.129099 |
| Human coronavirus 229E | 0.196892 | 0.597574 | 0.083607 | 0.121927 | 0.227798 | 0.552015 | 0.084688 | 0.135499 |
| Human coronavirus NL63 | 0.190087 | 0.712752 | 0.034993 | 0.062168 | 0.205129 | 0.677292 | 0.036428 | 0.081151 |
| Lucheng Rn rat coronavirus | 0.184384 | 0.557929 | 0.083031 | 0.174656 | 0.193101 | 0.527395 | 0.094169 | 0.185335 |
| Mink coronavirus | 0.228678 | 0.546826 | 0.102932 | 0.121563 | 0.249230 | 0.535591 | 0.099763 | 0.115417 |
| NL63-related Bat coronavirus | 0.189823 | 0.577504 | 0.079057 | 0.153616 | 0.210809 | 0.551366 | 0.075071 | 0.162754 |
| Porcine epidemic diarrhea virus | 0.199831 | 0.522930 | 0.096686 | 0.180553 | 0.191929 | 0.514827 | 0.105637 | 0.187607 |
| Rousettus bat coronavirus HKU10 | 0.207802 | 0.592856 | 0.091049 | 0.108293 | 0.209272 | 0.557211 | 0.083908 | 0.149609 |
| Scotophilus bat coronavirus 512 | 0.201188 | 0.544340 | 0.098665 | 0.155807 | 0.203487 | 0.569018 | 0.082646 | 0.144849 |
| Swine enteric coronavirus | 0.240513 | 0.573303 | 0.070222 | 0.115962 | 0.257887 | 0.516850 | 0.086973 | 0.138291 |
| Transmissible gastroenteritis virus | 0.246219 | 0.576135 | 0.057783 | 0.119863 | 0.270000 | 0.524111 | 0.082884 | 0.123004 |
| Wencheng Sm shrew coronavirus | 0.264157 | 0.548777 | 0.091945 | 0.095121 | 0.250293 | 0.609173 | 0.086062 | 0.054471 |
| **Gammacoronaviruses** | A4f | U4f | G4f | C4f | A4f | U4f | G4f | C4f |
| Avian infectious bronchitis virus | 0.280215 | 0.514909 | 0.105775 | 0.099101 | 0.272677 | 0.507519 | 0.096646 | 0.123157 |
| Beluga whale coronavirus | 0.251623 | 0.483031 | 0.137046 | 0.128301 | 0.258106 | 0.472517 | 0.137313 | 0.132064 |
| **Deltacoronaviruses** | A4f | U4f | G4f | C4f | A4f | U4f | G4f | C4f |
| Bulbul coronavirus HKU11 | 0.271396 | 0.538218 | 0.074722 | 0.115664 | 0.273612 | 0.512611 | 0.079387 | 0.134389 |
| Common-moorhen coronavirus | 0.280305 | 0.604411 | 0.059284 | 0.056000 | 0.245981 | 0.611189 | 0.064450 | 0.078381 |
| Magpie-robin coronavirus HKU18 | 0.160454 | 0.413506 | 0.141714 | 0.284326 | 0.199450 | 0.391917 | 0.128339 | 0.280294 |
| Munia coronavirus HKU13 | 0.271725 | 0.430193 | 0.116227 | 0.181855 | 0.281493 | 0.422466 | 0.100486 | 0.195556 |
| Night-heron coronavirus HKU19 | 0.355703 | 0.449160 | 0.068657 | 0.126479 | 0.362609 | 0.398874 | 0.078126 | 0.160391 |
| Porcine coronavirus HKU15 | 0.241219 | 0.471506 | 0.105017 | 0.182258 | 0.267771 | 0.423215 | 0.101061 | 0.207953 |
| Sparrow coronavirus HKU17 | 0.227610 | 0.427294 | 0.111387 | 0.233708 | 0.251201 | 0.419277 | 0.109639 | 0.219882 |
| Thrush coronavirus HKU12 | 0.266167 | 0.564516 | 0.071252 | 0.098065 | 0.276805 | 0.503051 | 0.098791 | 0.121352 |
| White-eye coronavirus HKU16 | 0.270696 | 0.499818 | 0.072899 | 0.156586 | 0.255200 | 0.502311 | 0.08055 | 0.161940 |
| Wigeon coronavirus HKU20 | 0.270351 | 0.483598 | 0.111698 | 0.134354 | 0.259563 | 0.486631 | 0.106387 | 0.147419 |

### Nucleotide usage bias analysis

The nucleotide usage biases along the length of ORF1 from each virus were calculated in sliding windows 150 codons in length, with a step of a single codon. The “VVTAK SW” algorithm (chemres.bsmu.by) was used for the calculations. Among other indices, the algorithm calculates nucleotide usage in fourfold degenerated sites, in which all nucleotide mutations are synonymous. ORF1b in each sequence was “opened up” by the addition of “A” nucleotide into the “AAA” motif of the ribosome slippage sequence. The location of this motif has been determined with the help of GenBank description of each sequence. So, the length of complete ORF1 for SARS-CoV-2 is equal to 7097 codons. This includes ORF1a equal to 4406 codons (until the stop-codon), and the ORF1b of 2696 codons. Nucleotide usages in the fourfold degenerate sites of both the parts of ORF1 (the one before the “AAA” motif of the ribosome slippage sequence, and the one after that motif) were compared for all 49 viruses. Coefficients of correlations between ΔU4f and ΔA4f, between ΔU4f and ΔC4f, between ΔU4f and ΔG4f, as well as between ΔA4f and ΔC4f, were calculated.

### Mutational pressure analysis in SARS-CoV-2

Nucleotide sequences of the complete ORF1 of SARS-CoV-2 were retrieved with the help of NCBI BLAST algorithm. The reference genome of SARS-CoV-2 was used as a target. 400 full length sequences of ORF1 belonging to different isolates of SARS-CoV-2 from all over the globe were obtained from GenBank on the 15^th^ of April 2020. The numbers of sites with all possible types of nucleotide mutations in this alignment relative to the reference sequence were calculated. Similarly, the number of sites available for each type of nucleotide mutation was calculated in the reference sequence. The rate of nucleotide mutation of a certain type is equal to the number of sites with a given mutation divided by the total number of nucleotides available for this kind of mutation. For example, the rate of C to U mutations is equal to the number of sites with C to U mutations over the number of C residues. The rates of mutations have been compared with each other with the help of t-test.

The rates of synonymous and nonsynonymous mutations for C to U mutations were calculated. The sites in which C to U mutation is synonymous and those in which it is nonsynonymous in the reference ORF1 sequence of SARS-CoV-2 were determined. To calculate the rate of synonymous C to U transitions, the number of sites with (non)synonymous C to U mutations were divided by the number of sites in which C to U mutations are (non)synonymous. The rates of mutations for both parts of the ORF1 were calculated separately.

### Statistical Analysis

The Student’s t-test was used for calculating the statistical significance. P-values < 0.05 were considered as statistically significant. Coefficients of correlation have been calculated using MS Excel.

## Results

### Mutational pressure in SARS-CoV-2 ORF1

As on the 15^th^ of April 2020, a total of 400 full-length sequences of ORF1 belonging to different isolates of SARS-CoV-2 were available in GenBank. The first full-length genome of SARS-CoV-2 appeared in GenBank towards the end of December 2019, followed by the other sequences. Therefore, the first reported genome of the virus can be considered as the initial sequence, while the others can be regarded as its offspring that mutated during the ongoing pandemic. If each offspring sequence is compared with the initial one, the number of mutations may seem to be relatively small. However, when all those 400 sequences are considered, there are already 250 sites with mutated nucleotides (ignoring ambiguous results of sequencing) in this enormously long ORF that consists of 21227 nucleotides.

To find out the preferable direction of those nucleotide mutations, the numbers of sites with each type of nucleotide substitution were calculated and divided by the usage of a corresponding nucleotide in a reference sequence^15^. Calculated frequencies of nucleotide mutations were compared with each other. Since there is a ribosome slippery sequence in the ORF1, we performed calculations separately for two parts of ORF1: ORF1a and ORF1b, before and after the slippery site, respectively. There is a clear and strong mutational U-pressure in both parts of the ORF1 (Table 1). The most frequent type of nucleotide mutation in SARS-CoV-2 ORF1 during 4 months of the pandemic was cytosine to uracil transition (C to U). The rate of this mutation was found to be more than 6 times higher than the rate of an opposite U to C mutation in ORF1a and more than 4 times higher in ORF1b. The difference between the rate of C to U transitions and the rate of U to C transitions is significant (P < 0.05) for both parts of ORF1. Interestingly, the rate of C to U transitions is also significantly (1.7 times) higher in the ORF1a than in ORF1b. The most frequent cause of this kind of mutation is cytosine deamination, the product of which is uracil. This mutation may be spontaneous or enzymatic. In the latter case, RNA-editing enzymes from APOBEC family may be responsible for it^13^,^14^. These enzymes bind single-stranded viral RNA and deaminate cytosine residues. Spontaneous deamination occurs via oxidation of cytosine by free radicals^16^. The rate of this process is higher for single-stranded RNA than for double-stranded RNA^17^.

Since ORF1a is translated more frequently than ORF1b, the rate of C to U transitions should be higher in ORF1a. Indeed, the process of translation often ends near the ribosome slippery sequence at a stop codon if the ribosome fails to slip to the -1 reading frame^6^. Therefore, ORF1b is not unwound as frequently as ORF1a. The difference between the rates of G to A and A to G transitions was not found to be significant for both parts of ORF1. However, the rate of G to A transitions themselves is significantly higher in ORF1a than in ORF1b (Table 1). The rate of G to U transversions is significantly higher than the rate of U to G transversions in both parts of ORF1 (Table 1). The rate of G to U transversions is, of course, significantly lower than the rate of C to U transitions. However, both of these mutations do contribute to the mutational U-pressure in this viral gene. The rates of other transversions are rather low (Table 1). Their preferable directions can be determined only after some time when the virus acquires more mutations. Even now, after the first four months after the breaking of the interspecies barrier by this virus, it is evident that the most expected mutations in its ORF1 are C to U transitions.

### The rate of synonymous C to U mutations is higher than the rate of nonsynonymous ones in SARS-CoV-2 ORF1

The number of C to U mutations observed in SARS-CoV-2 ORF1 is enough to make conclusions about the type of natural selection. To make such conclusions, the numbers of sites with synonymous and nonsynonymous C to U mutations in each part of ORF1were calculated^18^. After that, we divided those numbers by the numbers of sites for synonymous and non-synonymous C to U mutations in corresponding parts of ORF1 of the reference SARS-CoV-2 genome. A comparison of those rates is given in Table 2. As evident from Table 2, the rate of synonymous mutations of C to U direction is significantly higher than the rate of nonsynonymous mutations of the same direction in both parts of ORF1. The rate of synonymous C to U mutations is similar in both parts of ORF1. In contrast, the rate of nonsynonymous C to U mutations is significantly higher in ORF1a than in ORF1b.

The number of sites for synonymous C to U mutations is 2.7 (ORF1a) and 2.5 (ORF1b) times lower than the number of sites for nonsynonymous C to U mutations. So, the increase of the rate of C to U mutations in the first part of ORF1 relative to its second part is caused by the increase of the rate of nonsynonymous mutations. The reason for this growth maybe both in the higher rate of their occurrence and in the weaker negative selection. Indeed, proteins that are cut from the second part of a polyprotein encoded by ORF1b (RNA-dependent-RNA polymerase and helicase, 3’-to-5’ exonuclease, endoRNAse, 2’-O-ribose methyltransferase) seem to be more conservative than proteins that are cut from its first part. An interesting fact is that the number of sites with synonymous C to U mutations is lower than the number of sites with nonsynonymous C to U mutations (32 vs. 51) for ORF1a, while for ORF1b it is vice versa (17 vs. 13). As usual, these numbers must be normalized by the number of sites available for those mutations before the comparison.

### Nucleotide usage biases in ORF1 of SARS-CoV-2, SARS-CoV, and MERS

Nucleotide usage biases in genes are formed during long-term process of fixation of certain nucleotide mutations in numerous generations^19^. As one can see in Figure 1a, the usage of uracil in fourfold degenerated sites of ORF1 from SARS-CoV-2 is quite high: 53.6±0.2% in the first part of ORF1 and 49.0±0.2% in the second part implying that the rates of C to U transitions and G to U transversions have been much higher than the rates of opposite mutations for a very long time in predecessors of the current virus. Nowadays, almost one-half of nucleotides in fourfold degenerated sites of SARS-CoV-2 ORF1 are uracil residues. Interestingly, in ORF1b, the usage of uracil is still significantly lower than in the first one. It means that the tendency observed during the mutagenesis of SARS-CoV-2 in human cells is the same as during its mutagenesis in cells of its former hosts.

**Figure 1.**
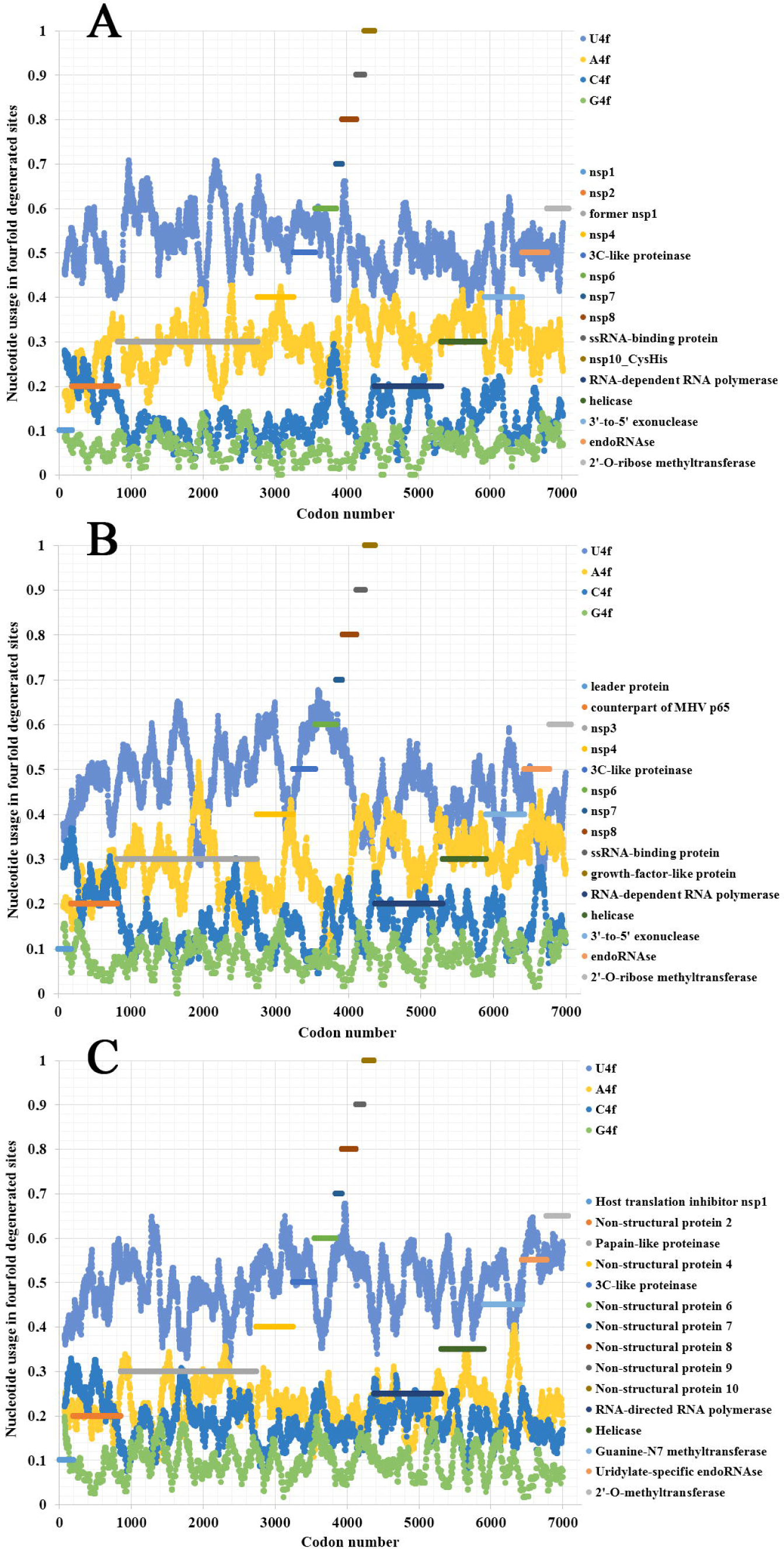
Nucleotide usage in fourfold degenerated sites along the length of ORF1 of a) SARS-CoV-2, b) SARS-CoV, c) Betacoronavirus England 1 (causative agent of MERS). The length of a sliding window is equal to 150 codons. Boarders of proteins that are cleaved from the long polypeptide are shown.

Adenine is the second most common/frequent nucleotide to occur in the fourfold degenerate sites (Figure 1a). This shows that C to U transitions are frequent in RNA minus strands of the virus as well. However, they are not as frequent as those in its RNA plus strands. This fact may be interpreted as the evidence that viral RNA plus strand is a target for APOBEC editing soon after the entry into the host cell. If an infection is successful, coronavirus suppresses expression of host proteins with its early protein nsp1^20^, and its RNA minus strands formed later are not edited by APOBEC. The level of A4f is significantly higher in the ORF1b than in ORF1a.

Both cytosine and guanine are extremely rare in fourfold degenerated sites of SARS-CoV-2 ORF1 (Figure 1a). However, there are still several areas with C4f level around 20% both in ORF1a and ORF1b.

In the first ORF of the SARS-CoV (responsible for the 2002-2003 SARS epidemic), the difference in nucleotide usage levels between two parts of ORF is even more noticeable than in SARS-CoV-2 (Figure 1b). In ORF1a, the level of U4f is 49.3±0.2%, while in ORF1b, it is 42.9±0.2%. The values of U4f for SARS-CoV ORF1 are significantly lower than those for SARS-CoV-2. As to the values of A4f, they are equal to 26.7±0.2% and 32.7±0.2% for SARS and 28.3±0.2% and 31.5±0.2% for SARS-CoV-2 ORF1 parts. So, the magnitude of change in A4f usage before and after the ribosome slippage site is higher for SARS-CoV than for SARS-CoV-2. It means that one may expect the highest rate of successful ribosome slippage for SARS-CoV-2 than for SARS-CoV.

Nucleotide usage biases in fourfold degenerated sites along the ORF1 of the MERS virus (namely, in Betacoronavirus England 1 strain) are shown in Figure 1c. The usage of U4f is distributed almost identically along the length of ORF1a (49.2±0.2%) and ORF1b (50.5±0.2%). A slight decrease of U4f in ORF1a is because of the area between codons #1400 and #2600, where U4f is lower than in other parts of the same ORF. The difference between A4f in ORF1a and ORF1b is insignificant (22.4±0.1% and 22.2±0.2%). There are areas with relatively elevated C4f usage in this ORF, while G4f is always somewhere near the point of 10%. Due to the absence of changes in nucleotide usage biases before and after the ribosome slippage sequence, one may speculate that this event (ribosome slippage) is almost always successful for the MERS virus. It is essential to check whether MERS virus is rather an exception or a rule among different coronaviruses.

### Nucleotide usage biases along the length of Alpha-, Beta-, Gamma-, and Deltacoronaviruses

Nucleotide usage biases along the length of ORF1 have been determined in 46 more completely sequenced coronaviruses. From the data given in Table 3, it can be concluded that there is a mutational U-pressure in all those known species. However, the average value of U4f in fourfold degenerated sites varies from 39 to 71%, while the average value of A4f varies from 16 to 40%. Once again, U4f is always significantly higher than A4f in all species of coronaviruses, in both parts of ORF1. If we take all 49 species of coronaviruses together, U4f is significantly higher in ORF1a than in ORF1b, while A4f and C4f are significantly lower in ORF1a than in ORF1b. The level of G4f is the same in both parts of ORF1. A similar trend is observed when only Alphacoronaviruses or Betacoronaviruses are considered. However, in Deltacoronaviruses, the differences in U4f and C4f in both parts are still significant, while the difference in A4f is not significant.

There are several exceptions from the rule among studied viruses, in which the value of U4f is significantly lower in ORF1a than in ORF1b. They are 3 out of 19 Alphacoronaviruses; 2 out of 18 Betacoronaviruses; 3 out of 10 Deltacoronaviruses (Table 3). There are at least two causes of the existence of those exceptions. At first, coronaviruses are prone to recombination with each other and with other viruses^21^. That is how some fragments of ORF1 may be exchanged during recombination, and a newly acquired fragment of ORF1 will possess an outstanding bias in fourfold degenerated sites. After a certain number of generations, nucleotide usage levels in fourfold degenerated sites will become almost identical through the whole length of ORF1a and ORF1b again. From this point of view, there may be such half-homogenized fragment in the ORF1a of MERS (Figure 1c). The “homogenization” of biases in twofold degenerated sites from third codon positions takes longer time than their “homogenization” in fourfold degenerated sites^22^. Biases in first and second codon positions will be “improved” after an even more extended period of time^23^.

Second, the quality of a regulatory element responsible for ribosome slippage should be different in different coronaviruses^6^. As one can see in Figure 2a, the difference in average values of U4f in two parts of ORF1 may reach more than 8%, or it may be equal to just 1%. Interestingly, there is a correlation between the difference in U4f and the difference in A4f between two parts of ORF1 (R = - 0.66). Indeed, the higher the difference in U4f, the higher (by module) the difference in A4f. The same relationships are there between the difference in U4f and the difference in C4f between two parts of ORF1 (R = - 0.72). The difference in A4f, however, shows no correlation on the difference in C4f (R = 0.11). It means that in some viruses, U4f in the first part of ORF1 is increased mostly because of the decrease of A4f, while in others, it is increased mostly because of the decrease of C4f. Both SARS-CoV-2 and SARS-CoV belong to the group in which U4f is growing mostly because of the decrease of A4f. The chance of success for ribosome slippage is one of the key factors of viral pathogenesis^24^. According to Figure 2a, SARS-CoV-2 should be able to synthesize more RNA-dependent-RNA-polymerase at the onset of infection than SARS-CoV.

**Figure 2.**
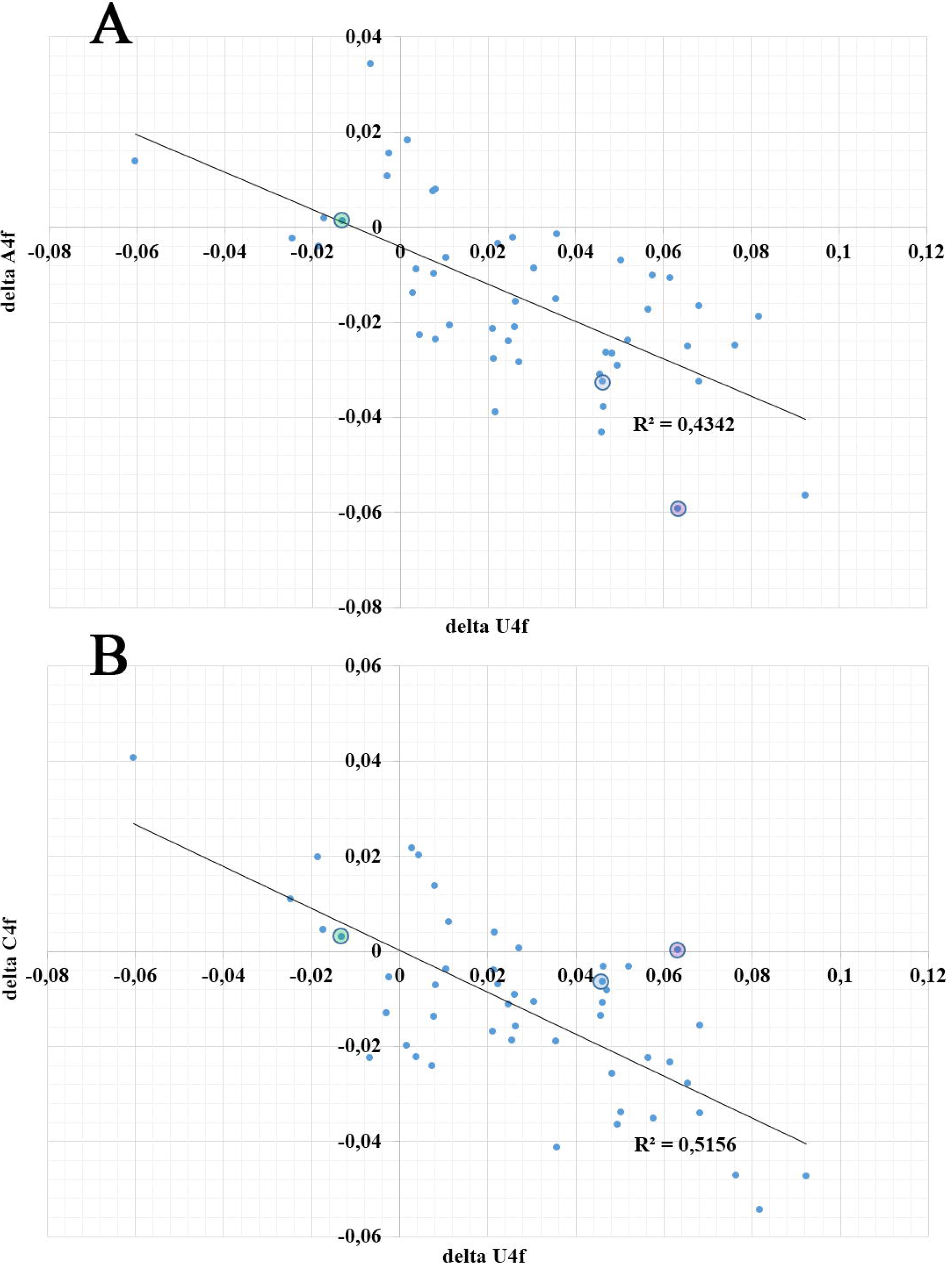
The dependence of the difference in A4f (a) and C4f (b) on the difference in U4f between ORF1a and ORF1b for 49 species of coronaviruses. Dots corresponding to SARS-CoV-2 are in blue circles, dots corresponding to SARS-CoV are in violet circles, dots corresponding to Betacoronavirus England 1 are in green circles.

## Discussion

Determination of the mutational pressure direction should be a starting point in any vaccine design study^22^. If the most frequent type of nucleotide mutation is known, one may try to choose “cold” spots for this mutation as targets for future vaccine development instead of “hot” spots. Based on our study, we suggest that fragments of RNA of SARS-CoV-2 that have a higher level of U in first and second codon positions and a higher level of C in synonymous sites for C to U mutations should be chosen for vaccine designing.

It has been shown that the genome of SARS-CoV-2 is subject to nucleotide usage bias towards A+U in its ORFs, including ORF1^25,26^. In this study, we examined the usage of U and A separately, since A=U and G=C parity rules do not work in these viruses. Nucleoside analogues have been suggested as anti-Covid-19 drugs^27^. Based on the results of this study, uracil analogues would be more effective in treating COVID-19 than analogues of cytosine and guanine nucleosides. Adenine analogues should be capable of effectively inhibiting the synthesis of RNA-minus strands. So they may also prove to be good therapeutic strategy but only during the early stages of infection. Though care should be taken that any nucleoside analogue used to treat Covid-19 wouldn’t be recognized by the proof-reading machinery of the virus^27^ or the drug will be rendered ineffective.

Sequences obtained from all over the world belong to active viruses that are under the control of negative selection, which can be confirmed by the fact that the rate of nonsynonymous mutations is less than the rate of synonymous ones. Indeed, the virus is not under strong immune pressure or the pressure of antiviral drugs at the moment. These stresses are known to cause diversifying (positive) selection of nonsynonymous mutations^28^. On the one hand, mutagenesis of C to U causes amino acid substitutions that can lead to quite drastic consequences for the fitness of a virus. Therefore, most of them are eliminated. On the other hand, mutations of C to U direction may help in immune evasion, since they are capable of destroying linear B-cell epitopes^29^.

Nsp1 is a non-structural protein present in Alpha- and Betacoronaviruses, but not in Gamma- and Deltacoronaviruses^20^. Nsp1 can inhibit the expression of host proteins.^30,31^ Likely, the expression of nonspecific antiviral enzymes belonging to APOBEC and ADAR families are also inhibited by nsp1. If most of the C to U mutations in ORF1a are caused by APOBEC editing of viral RNA during initial steps of infection, then it may be expected that there will be no difference in nucleotide usage levels in fourfold degenerated sites between both the parts of ORF1 in Gamma- and Deltacoronaviruses. But, as evident from Table 3, the above-mentioned differences still exist in most of the Gamma and Deltacoronaviruses, implying that a significant fraction of C to U mutations is not caused by APOBEC editing.

The existence of RNA exonuclease (nsp14) in genomes of coronaviruses may be the cause of the observed mutational U-pressure. If during the post-replicational proof-reading most of the deaminated and oxidized nucleotides are removed, the only product of deamination remaining is a canonical amine base, uracil. As inosine (product of adenine deamination) is a non-canonical amine base, it is removed during proof-reading, whereas, uracil (product of cytosine deamination) being a canonical amine base is not removed, as proof-reading activity effectively removes mismatched pairs of canonical nucleotides, i.e., pairs of canonical nucleotides with non-canonical ones, but it cannot recognize U that has already appeared in place of C on a matrix strand before the replication as a mutated nucleotide. Interestingly, some non-canonical amine bases, like 8-oxo-G (the product of guanine oxidation), can somehow still pass through the proofreading machinery, as several G to U transversions have been detected in this study.

Without proof-reading machinery, coronaviruses accumulate noncanonical nucleotides much better than in case when it is functional^32^. Indeed, 5-formyl-uracil is paired with adenine in the absence of nsp14 in SARS-CoV and MERS viruses, and it leads to the increase of U to C and A to G transitions since this non-canonical nucleotide can pair with G even better than with A^32^. So, in the absence of proof-reading, the overall rate of mutations becomes higher, while the bias in their rates becomes weaker.

Taken together, coronaviruses are well known for their low mutation rates achieved due to proof-reading during RNA replication^12^. However, they still cannot repair C to U transitions with this mechanism, as U is a canonical amine base. Moreover, C to U transitions occur before the replication, and the resulting U makes a correct pair with A during the complementary RNA strand synthesis. Therefore, mutational U-pressure is seen in all coronaviruses.

The effectiveness of frameshifting for ORF1 for different coronaviruses has been reported to be in the range of 20 – 45%.^6^. For the Mouse hepatitis virus A59, the rate of effective frameshifting is higher (from 48 to 70%)^33^. Indeed, ΔU4f for this virus is just 2.71%. For SARS-CoV, this rate has been reported as 17.5%,^6^, and its ΔU4f is 6.33%. In infectious bronchitis virus, the rate of successful frameshifting is 30 – 40%^7,34^, while ΔU4f is about 0.74%. However, roughly one-third of ORF1 at its 5’-end has surprisingly lower U4f values than the middle part, probably because of recombination events. For human coronavirus 229E the rate of successful frameshifting is about 20 – 30% ^35^, while ΔU4f is 4.56%. Hence, it may be speculated that highly efficient frameshifting leads to the decrease of ΔU4f, while low rate of effective frameshifting causes the increase of ΔU4f.

## Conclusions

In SARS-CoV-2 mutational U-pressure is associated with translation, as the rate of C to U transitions and the intensity of U-bias in fourfold degenerated sites are higher in ORF1a, situated before the ribosome slippery sequence, than in the less frequently translated ORF1b which is situated after the slippery sequence. U-pressure observed in all the examined sequences of ORF1 from different species of coronaviruses is a consequence of proof-reading.

## Author’s contribution

VVK, RG: conception and design. VVK, TAK: data acquisition, analysis, interpretation of data, and writing of the manuscript. VVK, RG, TAK, SAN, PVV, SKK review and writing of manuscript.

## References

1. Situation reports. https://www.who.int/emergencies/diseases/novel-coronavirus-2019/situation-reports .

2. Perlman, S. & Netland, J. Coronaviruses post-SARS: update on replication and pathogenesis. Nat. Rev. Microbiol. 7, 439–50 (2009).

3. Lauber, C. et al. Mesoniviridae: a proposed new family in the order Nidovirales formed by a single species of mosquito-borne viruses. Arch. Virol. 157, 1623–8 (2012).

4. Snijder, E. J., Decroly, E. & Ziebuhr, J. The Nonstructural Proteins Directing Coronavirus RNA Synthesis and Processing. in Advances in Virus Research vol. 96 59–126 (Academic Press Inc., 2016).

5. Nga, P. T. et al. Discovery of the first insect nidovirus, a missing evolutionary link in the emergence of the largest RNA virus genomes. PLoS Pathog. 7, e1002215 (2011).

6. Baranov, P. V et al. Programmed ribosomal frameshifting in decoding the SARS-CoV genome. Virology 332, 498–510 (2005).

7. Brierley, I., Digard, P. & Inglis, S. C. Characterization of an efficient coronavirus ribosomal frameshifting signal: requirement for an RNA pseudoknot. Cell 57, 537–47 (1989).

8. Ziebuhr, J., Snijder, E. J. & Gorbalenya, A. E. Virus-encoded proteinases and proteolytic processing in the Nidovirales. J. Gen. Virol. 81, 853–79 (2000).

9. Bass, B. L. RNA Editing by Adenosine Deaminases That Act on RNA. Annu. Rev. Biochem. 71, 817–846 (2002).

10. Sanjuan, R., Nebot, M. R., Chirico, N., Mansky, L. M. & Belshaw, R. Viral Mutation Rates. J. Virol. 84, 9733–9748 (2010).

11. Khrustalev, V. V., Khrustaleva, T. A., Sharma, N. & Giri, R. Mutational pressure in Zika virus: Local ADAR-editing areas associated with pauses in translation and replication. Front. Cell. Infect. Microbiol. 7, (2017).

12. Bouvet, M. et al. RNA 3′-end mismatch excision by the severe acute respiratory syndrome coronavirus nonstructural protein nsp10/nsp14 exoribonuclease complex. Proc. Natl. Acad. Sci. U. S. A. 109, 9372–9377 (2012).

13. Sharma, S. & Baysal, B. E. Stem-loop structure preference for site-specific RNA editing by APOBEC3A and APOBEC3G. PeerJ 2017, (2017).

14. Smith, H. C. RNA binding to APOBEC deaminases; Not simply a substrate for C to U editing. RNA Biology vol. 14 1153–1165 (2017).

15. Khrustalev, V. V., Khrustalerva, T. A., Poboinev, V. V. & Yurchenko, K. V. Mutational pressure and natural selection in epidermal growth factor receptor gene during germline and somatic mutagenesis in cancer cells. Mutat. Res. - Fundam. Mol. Mech. Mutagen. 815, 1–9 (2019).

16. Gros, L., Saparbaev, M. K. & Laval, J. Enzymology of the repair of free radicals-induced DNA damage. Oncogene vol. 21 8905–8925 (2002).

17. Hendriks, G., Jansen, J. G., Mullenders, L. H. F. & De Wind, N. Transcription and replication: Far relatives make uneasy bedfellows. Cell Cycle vol. 9 2300–2304 (2010).

18. Chi, P. B., Chattopadhyay, S., Lemey, P., Sokurenko, E. V. & Minin, V. N. Synonymous and nonsynonymous distances help untangle convergent evolution and recombination. Stat. Appl. Genet. Mol. Biol. 14, 375–389 (2015).

19. Sueoka, N. Wide intra-genomic G+C heterogeneity in human and chicken is mainly due to strand-symmetric directional mutation pressures: dGTP-oxidation and symmetric cytosine-deamination hypotheses. in Gene vol. 300 141–154 (2002).

20. Shen, Z. et al. A conserved region of nonstructural protein 1 from alphacoronaviruses inhibits host gene expression and is critical for viral virulence. J. Biol. Chem. 294, 13606–13618 (2019).

21. Su, S. et al. Epidemiology, Genetic Recombination, and Pathogenesis of Coronaviruses. Trends in Microbiology vol. 24 490–502 (2016).

22. Khrustalev, V. V. et al. The history of mutational pressure changes during the evolution of adeno-associated viruses: A message to gene therapy and DNA-vaccine vectors designers. Infect. Genet. Evol. 77, (2020).

23. Khrustalev, V. V., Arjomandzadegan, M., Barkovsky, E. V. & Titov, L. P. Low rates of synonymous mutations in sequences of Mycobacterium tuberculosis GyrA and KatG genes. Tuberculosis 92, 333–344 (2012).

24. Plant, E. P., Sims, A. C., Baric, R. S., Dinman, J. D. & Taylor, D. R. Altering SARS coronavirus frameshift efficiency affects genomic and subgenomic RNA production. Viruses 5, 279–294 (2013).

25. Kandeel, M., Ibrahim, A., Fayez, M. & Al-Nazawi, M. From SARS and MERS CoVs to SARS-CoV-2: Moving toward more biased codon usage in viral structural and nonstructural genes. J. Med. Virol. (2020) doi:10.1002/jmv.25754.

26. Sheikh, A., Al-Taher, A., Al-Nazawi, M., Al-Mubarak, A. I. & Kandeel, M. Analysis of preferred codon usage in the coronavirus N genes and their implications for genome evolution and vaccine design. J. Virol. Methods 277, (2020).

27. Agostini, M. L. et al. Small-Molecule Antiviral β-D-N 4 -Hydroxycytidine Inhibits a Proofreading-Intact Coronavirus with a High Genetic Barrier to Resistance . J. Virol. 93, (2019).

28. Nijmeijer, B. M. & Geijtenbeek, T. B. H. Negative and positive selection pressure during sexual transmission of transmitted founder HIV-1. Frontiers in Immunology vol. 10 (2019).

29. Khrustalev, V. V. Levels of HIV1 gp120 3D B-cell epitopes mutability and variability: Searching for possible vaccine epitopes. Immunol. Invest. 39, 551–569 (2010).

30. Tanaka, T., Kamitani, W., DeDiego, M. L., Enjuanes, L. & Matsuura, Y. Severe Acute Respiratory Syndrome Coronavirus nsp1 Facilitates Efficient Propagation in Cells through a Specific Translational Shutoff of Host mRNA. J. Virol. 86, 11128–11137 (2012).

31. Huang, C. et al. Alphacoronavirus Transmissible Gastroenteritis Virus nsp1 Protein Suppresses Protein Translation in Mammalian Cells and in Cell-Free HeLa Cell Extracts but Not in Rabbit Reticulocyte Lysate. J. Virol. 85, 638–643 (2011).

32. Smith, E. C., Blanc, H., Vignuzzi, M. & Denison, M. R. Coronaviruses Lacking Exoribonuclease Activity Are Susceptible to Lethal Mutagenesis: Evidence for Proofreading and Potential Therapeutics. PLoS Pathog. 9, (2013).

33. Irigoyen, N. et al. High-Resolution Analysis of Coronavirus Gene Expression by RNA Sequencing and Ribosome Profiling. PLoS Pathog. 12, (2016).

34. Dinan, A. M. et al. Comparative Analysis of Gene Expression in Virulent and Attenuated Strains of Infectious Bronchitis Virus at Subcodon Resolution. J. Virol. 93, (2019).

35. Herald, J. & Siddell, S. G. An ‘elaborated’ pseudoknot is required for high frequency frameshifting during translation of HCV 229E polymerase mRNA. Nucleic Acids Research vol. 21 (1993).

